# Monodisperse LNPs - from Efficient Microfluidic Production and Loading to in Vitro Testing

**DOI:** 10.64898/2025.12.23.696212

**Authors:** Ebrahim Taiedinejad, Alshaimaa Abdellatif, Victor Krajka, Andreas Dietzel

**Author notes:** **Corresponding Author:** Ebrahim Taiedinejad, Andreas Dietzel.

## Abstract

Carrier nanoparticles facilitate the encapsulation of drug or mRNA molecules thereby enhancing their bioavailability. Microfluidic mixers provide a unique environment for the precise and continuous generation of nanoparticles by antisolvent precipitation. A major challenge is to understand the influence of microfluidic channel designs and geometries on the continuous production of small, uniform lipid nanoparticles (LNPs) and to identify conditions that ensure effective and controllable mixing of aqueous and organic phases in laminar flows. Another important challenge is that sufficient quantities for preclinical and clinical studies must be produced within a reasonable period of time. With this dual objective, different versions of a low aspect ratio laminar mixer (LARLM) were produced using two-photon polymerization (2PP). In the LARLM the organic phase forms a thin layer a few micrometers with a uniform velocity distribution in the center of the channel, surrounded by the aqueous phase. This concept has three major advantages: Firstly, it keeps all particles centralized in the channel, thus preventing contamination during prolonged particle generation. Secondly, diffusive mixing in the thin central stream occurs very quickly, and thirdly, the growing nanoparticles move at a homogeneous speed, which enables inline measurement of the particles. In systematic experiments with design versions of varied channel dimensions the operational parameters such as lipid concentrations and flow rates and the capability to produce LNPs with desired properties and loading capacities were explored. An interfacial dispersion model (IDM) could explain the surprising reduction of particle sizes with increased productivity. The latter allowed us to produce nanoparticles in the range of 50 nm to 180 nm (with 0.02 < PDI < 0.1) under stable conditions with a productivity of around one liter of LNP suspensions every three hours. Such performance has never been reached before with microfluidics. Moreover, LNPs loaded with coumarine-6 and various drugs were produced in the LARLM. Moreover, in-vitro experiments could confirm an improved bioavailability of coumarin-6 in cell culture experiments when loaded in LNPs by the LARLM. These results highlight the unique capabilities of LARLM devices and their potential to support nanoparticle formulation studies including preclinical and in further developments also clinical studies as required for approval as a marketable medicine.

## 1. Introduction

In the evolving landscape of pharmacotherapy, lipid nanoparticles (LNPs) have emerged as pivotal vehicles for designing efficient drug delivery. The significance of LNPs has been underscored by their role in the rapid development and deployment of mRNA vaccines, such as those for COVID-19. This success has catalyzed a broader trend towards mRNA therapies in various medical fields, including oncology and infectious diseases. The ability of LNPs to efficiently encapsulate, protect and deliver nucleic acids as well as to facilitate the uptake by target cells positions them as a preferred platform for innovative therapies, leading to increased research and clinical trials focused on their application ^1^.

The size of lipid nanoparticles has a major impact on their cellular uptake, which is mainly through endocytosis, where substances become engulfed by the phospholipid bilayer of the cell membrane. For example, similar to viruses such as influenza (80 nm) and retroviruses (120 nm), RNA-lipid nanoparticle (RNA-LNP) complexes of about 100 nm are mainly taken up by receptor-mediated endocytosis. Furthermore, optimal particle properties, including size and surface chemistry, vary between species, with nonhuman primates preferring smaller RNA LNP sizes of 50 -60 nm, while rodents prefer 70-80 nm.^2–5^ LNPs offer a multifaceted approach to advanced drug delivery. With regards to poorly water-soluble drugs, nanoparticle formulations contribute to enhanced dissolution and absorption, consequently improving bioavailability. Furthermore, through targeted drug delivery, nanocarriers act with greater effectiveness and decrease side effects. Simultaneously nanoparticles can exercise control over the release rate and minimize the degradation rate ^*6*^. Narrow size distributions, often measured by the polydispersity index (PDI), ensure kinetic stability of particles by mitigating Ostwald ripening process where larger particles grow at the expense of smaller ones due to differences in solubility ^7^. Fortunately, the rapid and uniform mixing capabilities of microfluidic tools play a key role in meeting these essential criteria. Homogenous and rapid mixing promotes an even and swift supersaturation throughout the system. The strategic manipulation of key parameters, including microchannel design as well as flow rates of injected phases, can control the mixing. This, in turn, triggers a prompt and uniform nucleation process, fostering small particles with a narrow size distribution^8^.

There exist numerous techniques for manufacturing nanoparticles. Microfluidic systems, employing the bottom-up approach based on the principle of antisolvent-nanoprecipitation, are advancing to establish their suitability for clinical application^9^. Microfluidic systems offer a solution to specific challenges encountered in top-down approaches (e.g. high-pressure homogenization, ultrasonication), steering clear of issues like high thermal energy input and shear stress^10,11,12^. The merging of aqueous and organic phases takes place by junctions or 3D nozzles in the microfluidic channels, initiating the diffusion of organic solvents into the aqueous phase^6,13^. This diffusion results in the supersaturation of lipids within the aqueous medium. Above a specific supersaturation threshold, nucleation is prompted. As the solute concentration descends below this critical threshold, existing nuclei undergo growth through either condensation or coagulation, without the formation of new nuclei. This process ultimately culminates in the formation of LNPs^14^. Microfluidic mixers promise exceptional control over the PDI. This precision is crucial for optimizing pharmacokinetics and ensuring effective cellular uptake, which directly impacts the therapeutic efficacy ^15^. The laminar flow characteristics inherent in microfluidic systems ensure consistent mixing conditions. This results in highly reproducible formulations, which is vital for maintaining batch-to-batch consistency in pharmaceutical applications^16,17^.

Despite all the positive aspects of microfluidics, there are two weaknesses that typically hinder their suitability to support clinical applications. One is low throughput and the other is fouling leading to insufficient stability in long term operation. Fouling is characterized by the deposition of material on the channel walls (particularly noticeable in passive micro fluidizers with a high surface-to-volume ratio), which can lead to shifts in the streamlines, clogging of the system as well as contaminations by material redetaching from channel walls and thus to an impairment of the continuity of the process. While various strategies such as ultrasound, high shear stress, inlet filters, specific channel materials and tailored solution compositions have been proposed to mitigate fouling in microfluidic systems, the search for a micro fluidizer suitable to support long-term stable production as required for clinical applications continues.

Recently, we have developed the concept of the Low Aspect Ratio Lamination Mixer (LARLM) ^17^-This microfluidic system with a sophisticated internal structure that facilitates 3D hydrodynamic flow focusing, was made possible by two-photon polymerisation (2PP). The LARLM ensures exclusivelydiffusive-driven mixing of organic and aqueous phases under laminar conditions. With a flat rectangular injection nozzle, which offers a larger surface area compared to round nozzles, an almost one-dimensional flow profile is created. The LARLM not only allows in-situ DLS feedback by concentrating the produced nanoparticles in the center of the one-dimensional flow, which is critical for convection compensation in the dynamic light scattering (DLS) speckle images, but also successfully prevents fouling. Moreover, it showed high productivity with tunable particle size.

However, systematic experiments with different channel dimensions, operating parameters such as lipid concentrations and flow rates are still lacking for the ability to produce LNPs with the desired properties and loading capacities. These should address the question whether LARLM devices enable the required productivity and potential to support nanoparticle formulation studies including preclinical and also clinical studies as required for approval as a marketable medicine.

## 2. Methods and materials

### 2.1 CFD simulations

CFD simulations of flow behavior in different LARLM designs were conducted using COMSOL Multiphysics 6.1 under a steady-state, single-phase flow conditions to analyze velocity distributions and details of hydrodynamic focusing. The incompressible Navier-Stokes equations were solved using the laminar flow module, with boundary conditions set to match experimental volumetric flow rates of the aqueous phase *Q*_*aqueous*_and the organic phase *Q*_*organic*_.

### 2.2 Femtosecond laser preparation of substrates and 2PP

1.1 mm thick 4” wafers of borosilicate glass (Borofloat® 33 from Schott, Mainz, Germany) served as the basis for the microfluidic chip. To enable good adhesion of 2PP printed microchannels and to create fluidic in- and outlets the glass was roughened together with alignment marks, and locally perforated using a femtosecond laser micro ablation system (microSTRUCT c, 3D Micromac AG, Chemnitz, Germany) in which a galvanometer scanner (Scanlab RTC5, Puchheim, Germany) moved the laser spot along parallel lines at a speed of 2000 mm/s. The structure was rotated 45° after one full pass and the laser focus was moved 50 μm deeper into the substrate on every fourth pass. The entire wafer was finally diced into separate 19 × 9.5 mm^2^ chips.

A 2PP system (Photonic Professional GT2+, Nanoscribe GmbH, Eggenstein-Leopoldshafen, Germany) was used to print microchannel architectures on individual chips. This technique exposed voxels with sub-micron lateral dimensions in a light-curing resist (IPS negative-tone photoresist, Nanoscribe GmbH, Eggenstein-Leopoldshafen, Germany) with a refractive index of 1.515 and a Young’s modulus of 5.1 GPa. A laser with a repetition rate of 80 MHz and a power of 63 mW was used to produce light pulses shorter than 90 fs at 780 nm. The laser was focused with a microscope objective lens (Zeiss LCl “Plan-Neofluar” 25×/0.8 Imm Korr Ph2) with a numerical aperture of 0.8 and a 25x magnification. Following laser exposure, the unpolymerized photoresist underwent removal in a developer bath (mr-Dev 600, micro resist technology GmbH, Berlin, Germany). Any remaining resist residues within the microchannels were removed by suction through the channel outlet. The dried prints underwent curing on a hotplate at 190°Cfor 10 minutes. The resultant transparent structures exhibited hydrophobic surface characteristics.

### 2.3 Microscopy

Scanning electron microscopy (Phenom XL, Phenom-World from Thermo Fisher, USA) was used to examine the shape and surface morphology of the channels. The flow conditions were investigated by light microscopy (Zeiss Axio Vert.A1, Zeiss, Oberkochen, Germany) using colored solutions. 2 mg/mL methylene blue (Sigma Aldrich, Steinheim, Germany) dissolved in absolute ethanol (Fisher Scientific, Loughborough, United Kingdom), representing the organic phase, was injected through the nozzle while deionized water was flushed through the main channel. To follow the flow of LNPs loaded with coumarin-6 (Assay > 99%, Sigma Aldrich, Saint Louis, MO 63103, USA) with fluorescent emission at λ= 505 nm in ethanol, a fluorescence microscope (IX73, Olympus, Tokyo, Japan) was used after a solution containing ethanol and 0.25 mg/mL coumarin-6 was injected into the main channel for 2 minutes so that the channel edges also became visible.

### 2.4 Offline DLS measurement

The nanodispersions were analyzed by measuring their Z-average and PDI using DLS (Nanoseries ZS Zetasizer -Malvern Instruments, UK). The Z-average reflects the hydrodynamic diameter of the particles, weighted by intensity. The suspensions obtained with varied flow rate ratios were diluted so that the solvent concentration in all samples in the cuvettes became negligible. A 1 ml sample was placed in a polystyrene cuvette (Omnilab GmbH, Bremen, Germany) which was equilibrated at 25 °C for 5 min before three measurements were taken at a backscatter angle of 173° for 120 s each. The calculations for the particle size parameterswere based on a dynamicviscosity of 0.8872 mPas and a refractive index of 1.330 for water at 25 °C.

### 2.5 LARLM design variants

Four variants of the LARLM were produced as shown in figures 1a-d. LARLM Ⓐ and LARLM Ⓑ have the same main channel crosssection of 750µm× 360 µm, but differ in injection nozzle dimensions. LARLM Ⓐ has a nozzle outlet of 516 µm × 10 µm, while the LARLM Ⓑ has a wider nozzle outlet of 516 µm × 30 µm. LARLM Ⓒ exhibits a more compact design with a main channel cross section of 750 µm × 200 µm and nozzle outlet dimensions of 516 µm × 30 µm (as LARLM Ⓑ). In LARLM Ⓓ the height is reduced resulting in a main channel cross section of 500 µm × 360 µm and nozzle outlet dimensions of 276 µm × 30 µm. The distance between the nozzle outlet and the end (outlet) of the main channel was 9.5 mm in all variants (see figure 1e). Figure 1f shows the inside of the injection nozzles with cascaded obstacles that ensure homogeneous injection of the organic phase into the aqueous phase, which is crucial for the homogeneous thickness of the phase injected through the nozzle.

**Figure 1.**
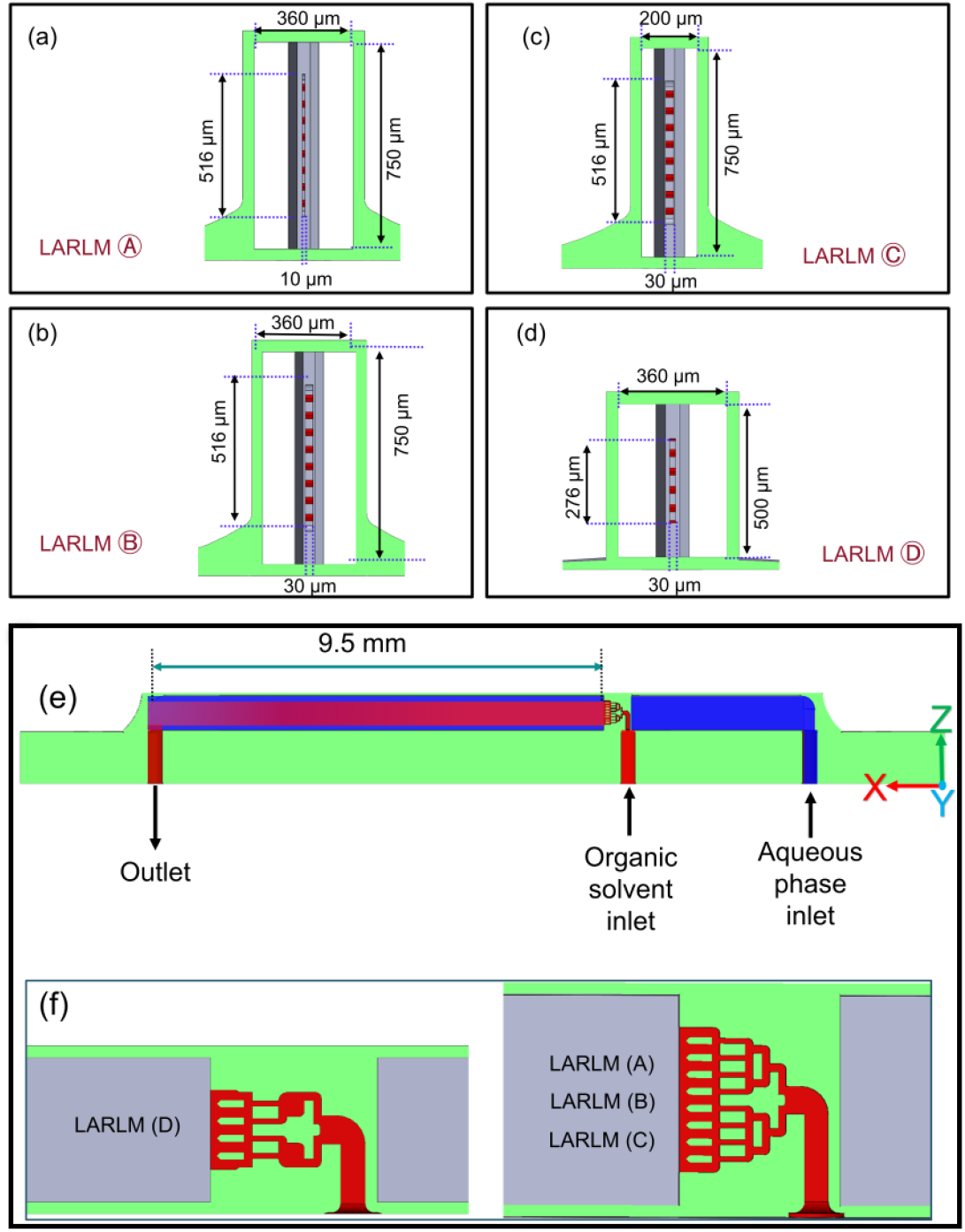
Schematic representations of different LARLM designs, illustrating variations in main channel cross-sections and nozzle dimensions: (a) LARLM Ⓐ, (b) LARLM Ⓑ, (c) LARLM Ⓒ, and (d) LARLM Ⓓ. The green regions indicate the structures that delimit the microchannels, while the gray and red areas represent the fluid-carrying microchannels. (e) Cross section in the x-z plane showing the organic phase injection and the positioning of the aqueous phase inlet and the outlet. (f) Interior of the LARLM nozzles with flow cascaded obstacles for flow homogenization, where the right section displays LARLM Ⓐ, LARLM Ⓑ, and LARLM Ⓒ, while the left section represents LARLM Ⓓ.

Figure 2a shows scanning electron microscope (SEM) images of the 2PP printed LARM chips confirming the cross-sectional geometries and figure 2b is a view of a partially opened chip to illustrate the injection of two phases. Moreover, filters with 10 µm pore diameter are shown which were integrated in the inlets to improve process stability. The finished chip, printed on a glass substrate is shown in figure 2d.

**Figure 2.**
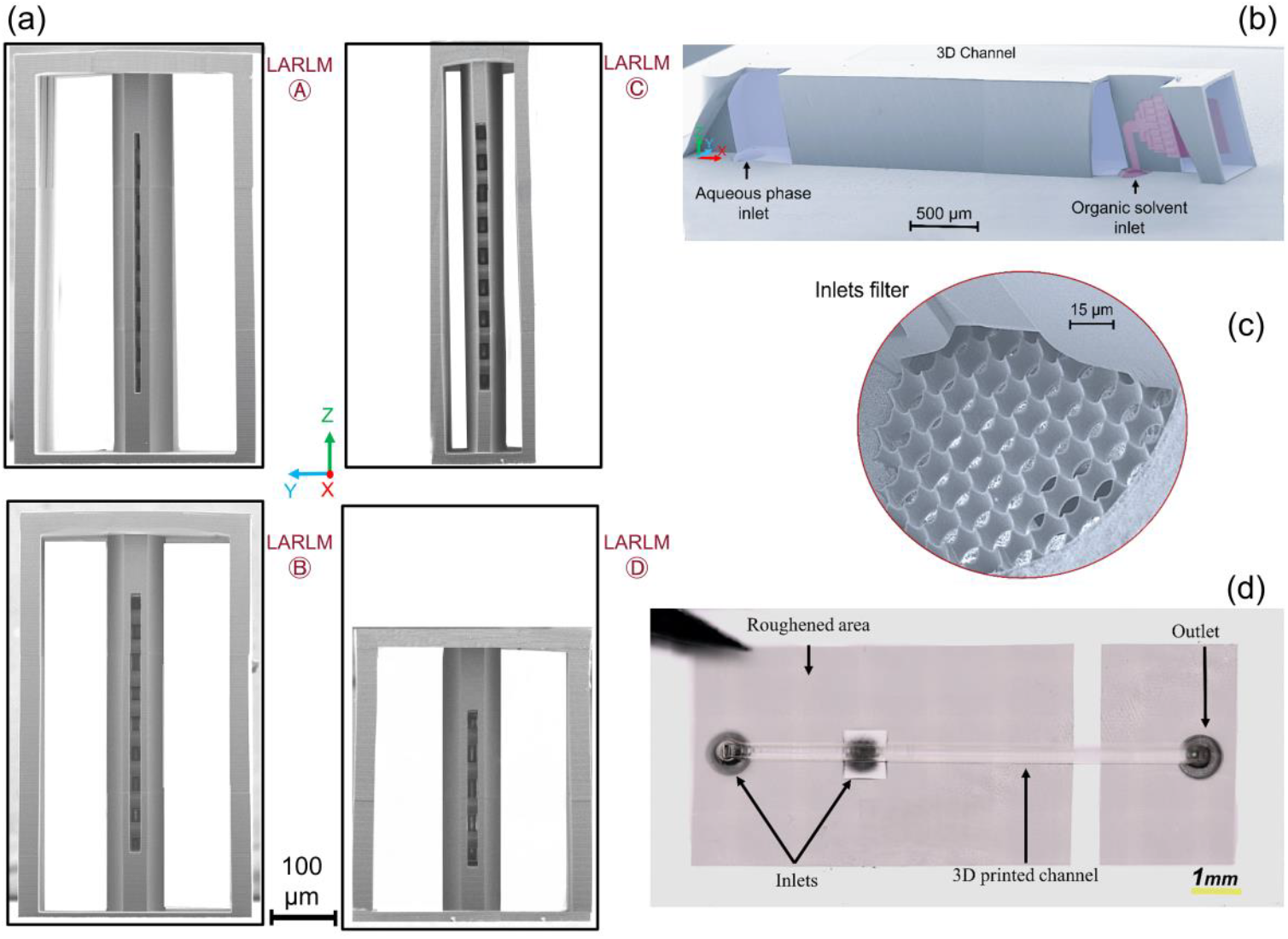
(a) SEM micrographs of LARLM Ⓐ, Ⓑ, Ⓒ and Ⓓ cross sections with injection nozzles and b) of the LARLM Ⓑ with partial openings to show nozzle interior with digital coloring illustrating the liquid phases (red: organic, blue: aqueous) as well as (c) filter printed into both inlets. (d) Photograph of a finished LARLM chip.

### 2.6 Experimental setup for LNP production and drug loading

As shown in figure 3, the chips were positioned between two aluminum mounting brackets to connect to the pumps. To ensure a secure seal, O-rings were placed between the holes in the aluminum brackets and those in the glass substrate. Once the brackets were tightened with screws, this design effectively prevented leakage. The assembly was connected to the syringes using PTFE tubing and flangeless screws (Techlab GmbH Braunschweig, Germany) for fluid delivery. Flow rates *Q*_*aqueous*_ and *Q*_*organic*_ (in loading experiments coumarin-6 and drugs were added to the organic solution) were precisely controlled using Nemesys Base120^+^ low pressure syringe pump modules (Cetoni GmbH, Korbussen, Germany) fitted with 2.5 mL glass syringes (Innovative Labor Systeme, Stützerbach, Germany).

**Figure 3.**
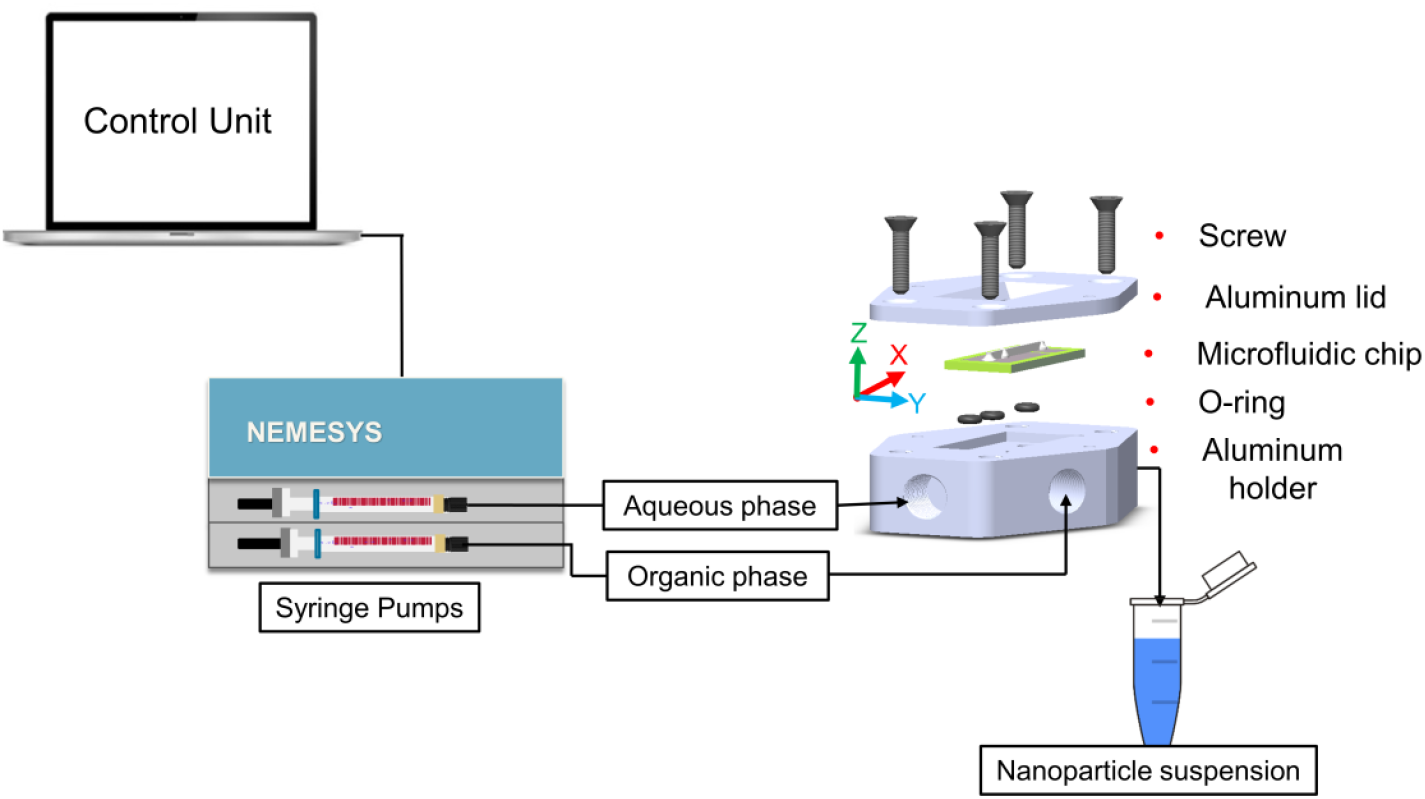
Sketch illustrating the experimental setup for injecting organic and aqueous phases into the LARLM channels together with an exploded view of the mounting bracket assembly with the chip.

### 2.7 Preparation of reagent

For the organic phase 5 mg mL^−1^ castor oil (Henry Lamotte Oils, Bremen, Germany) adhering to Ph. Eur. standards was dissolved in HPLC grade ethanol (Fisher Scientific, Loughborough, United Kingdom) and 2.5 mg mL^−1^ non-ionic surfactant polysorbate 80 (Sigma Aldrich, Steinheim, Germany) meeting Ph. Eur. specifications was added. For the aqueous phase purified water was obtained via reverse osmosis and sterile filtration using water purifier (Astacus from membraPure, Hennigsdorf, Germany). For the loading experiments coumarin-6 was added as a synthetic derivative. While some biological activities are associated with coumarin-6, such as antioxidant effects, its primary role lies in observing bioavailability through fluorescence.^18 19^ Moreover LNPs loaded with simvastatin (≥97% HPLC grade, Sigma Aldrich) and Diclofenac Sodium Salt (Sigma Aldrich, Steinheim, Germany) in order to see the drug effects on particle size and PDI. All active ingredients were added in concentrations of 0.25 mg mL^-1^ and 1 mg mL^-1^ in the organic phase. Prior to loading the syringes, both the aqueous and organic phases were filtered using 0.2 μm filters (Puradise 25 TF WhatmanTM, 0.2μm polytetrafluoroethylene, GE Healthcare UK).

### 2.8 Encapsulation efficiency measurement

Samples (1 ml each) of both LNP-encapsulated and non-encapsulated coumarin-6 were subjected to centrifugation at 14,800 rpm for 45 minutes in a Fresco centrifuge (Thermo Scientific, GmbH). After centrifugation, the supernatant was carefully separated from the sedimented coumarin-6. The absorption of the supernatant was measured using a UV-Vis spectrophotometer (Cary 60, Agilent Technologies) after a calibration curve with known concentrations of coumarin-6 was established to determine *C*_*en*_, which is the concentration of (encapsulated) coumarin-6 in the supernatant. The percentage encapsulation efficiency EE% was calculated as

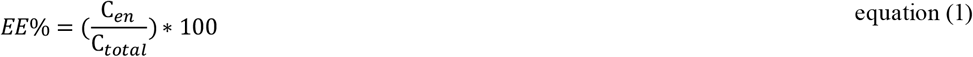

where *C*_*total*_ is the initial coumarin-6 concentration measured before centrifugation.

### 2.9 In vitro tests and fluorescence microscopic analysis

HUVECs (human umbilical vein endothelial cells, C-12203, PromoCell, Germany) at passages 3–5 were used to assess the uptake of coumarin-6-loaded LNPs. Cells were maintained in endothelial cell growth medium (C-22010, PromoCell) and cultured on gelatin-coated multiwell plates (07903, STEMCELL, Germany). For uptake experiments, cells were incubated for 30 min with LNP solutions (1:50 dilution in medium) prepared using LARLM Ⓑ at a constant flow rate ratio (*Q*_*aqueous*_ /*Q*_*organic*_ = 20) but varying total flow rates (*Q*_*aqueous*_ + *Q*_*organic*_ = 420, 4200, and 6300 µl/min). Microfluidic samples without lipid solutions served as controls. After incubation, cells were washed twice with medium. To visualize the plasma membrane, cells were stained with WGA-AF594 (2 µg ml^-1^, ABD-25509, Biomol GmbH, Germany) for 5 min before imaging. Fluorescence microscopy (IX73, Olympus, Tokyo, Japan) was performed at 30 min and 4 h after incubation, with fixed exposure times. LNP intensities were quantified using TrackMate^20^ with the Cellpose^21^ detector (Cytoplasm 2.0 model, cell diameter 100 pixels ≙ 33 µm). Cell segmentation was based on the WGA signal, and coumarin-6 signals were measured within segmented regions. Background levels were determined from untreated cells, and the median background intensity was subtracted from the coumarin-6-treated sample values.

## 3. Results and Discussion

Results of CFD simulations of convective transport (without taking transport by diffusion into account) in different LARLM designs at a total flow rate *Q*_*aqueous*_ + *Q*_*organic*_ = 420 μl min^-1^ and a flow rate ratio of *Q*_*aqueous*_ /*Q*_*organic*_= 20 are shown in figure 4a. Hydrodynamic focusing of the centrally injected organic phase by the co-flowing aqueous phase generates a thin sheet exhibiting only small spatial thickness variations. The velocity distributions of the organic phase are shown in x-z and y-z cross-sections, experimentally obtained phased is tributions in figures 4b-d are in agreement with the simulations. LARLM Ⓑ seems to produce a slightly more homogeneous velocity than LARLM Ⓐ and produces an injected sheet that extends slightly further in z-direction. Because the flow has to pass smaller cross sections, LARLM Ⓒ and LARLM Ⓓ exhibit increased velocities within the organic sheet layers.

**Figure 4.**
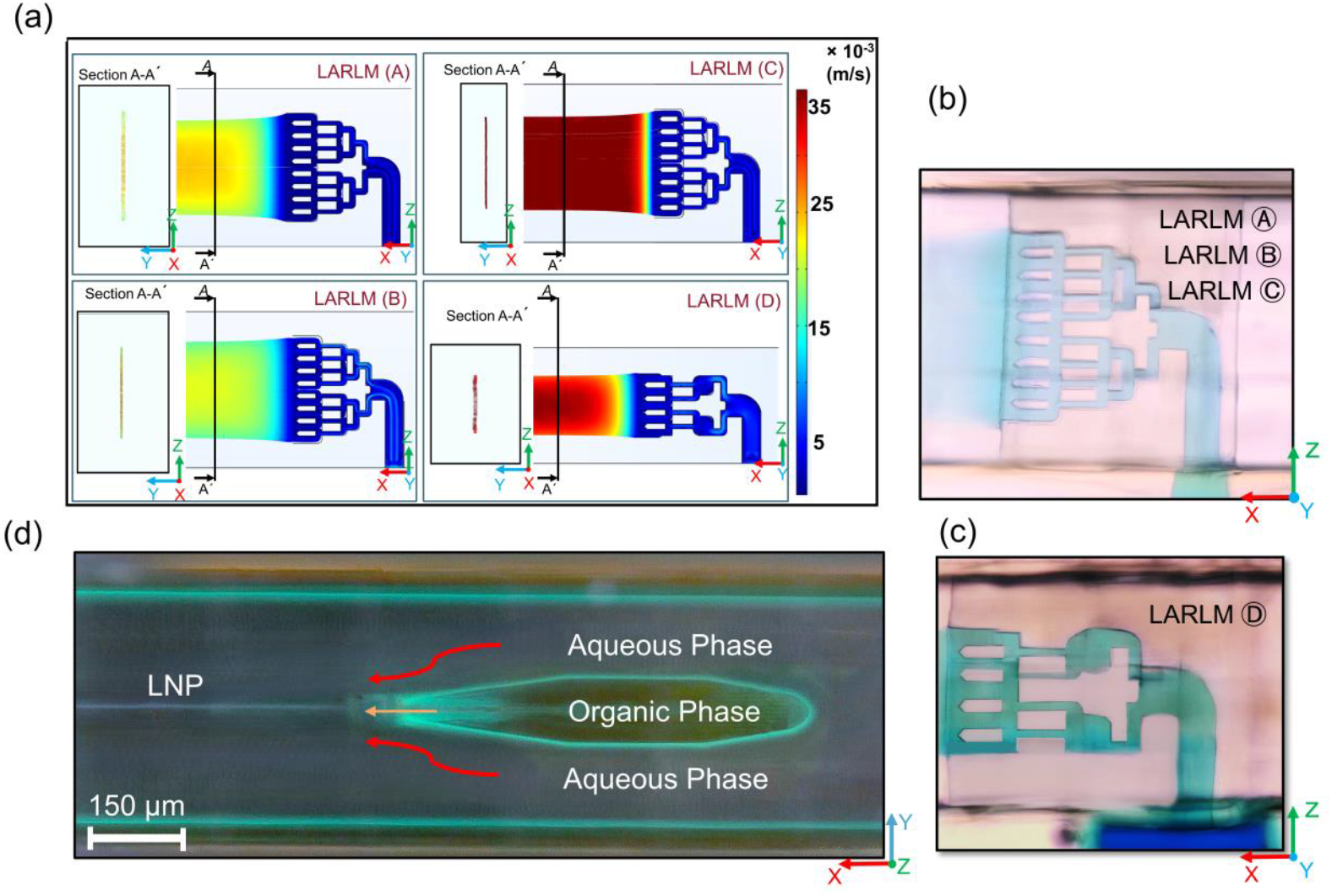
(a) Results of streamline simulations showing the injected organic phase in x-z and y-z cross sections in the four different LARLM designs as obtained with *Q*_*aqueous*_ + *Q*_*organic*_ = 420 μl min^-1^ and *Q*_*aqueous*_ /*Q*_*organic*_ = 20. The y-z cross sections were taken along A-A. The organic phase sheet thickness (in y-direction) has been enlarged threefold in the representation. (b) Microscopic view of the injected organic phase (green/blue color added) in side view as obtained almost identical for LARLM Ⓐ, LARLM Ⓑ, LARLM Ⓒ and (c) as obtained for LARLM Ⓓ. (d) Fluorescence microscopy image (x-y plane) of organic phase loaded with coumarine-6 moving in the channel center while channel walls have been stained with a mixture of absolute ethanol and 0.25 mg/ml coumarine-6.

From the simulations the thickness of the organic layer *d* for *Q*_*aqueous*_ /*Q*_*organic*_ = 20 was determined in all LARLM systems 9 mm downstream of the injection point where the flow has stabilized. Figure 5a shows that in all designs *d* does not depend on the total flow rate. In LARLM Ⓐ *d* is only marginally lower than in LARLM Ⓑ. This difference can be explained by the previously discussed slightly wider extension of the organic sheet in z-direction. LARLM Ⓒ produces a thinner sheet because the flow has to pass smaller cross section of the main channel. Figures 5b and 5c show the experimentally obtained nanoparticle sizes and PDI values at varied total flow rates. Reproducibility was confirmed through triplicate experiments across independently fabricated microfluidic channels. In agreement with previous studies with the LARLM the particle sizes do not significantly change up to *Q*_*aqueous*_ + *Q*_*organic*_= 2000 μL/min, while PDI values slightly increase.^16,17^ However, for even higher flow rates which have not been studied before a clear decrease in particle sizes is observed for all designs, despite the fact that *d* remains constant for all designs. The fact that LARLM Ⓒ produces the smallest particles is not a true advantage of this design, since this can easily be compensated by adjusting the flow rate ratio with a geometric ratio of approx. 170µm /330µm = 1.9 for the other designs. Switching from a 10 μm nozzle in LARLM Ⓐ to 30 μm nozzle in LARLM Ⓑ results in a slightly higher value of *d* obtained at 9 mm distance from the nozzle. Despite the initial assumption that this would decrease *t*_*diffusion*_ and reduce nanoparticle size and PDI, it has been found that this has the opposite effect. This observation together with the decrease of particle sizes at very high flow rates cannot be explained by the layer thickness *d* obtained after complete flow stabilization and must therefore result from not fully developed flow profiles in regions closer to the nozzle.

**Figure 5.**
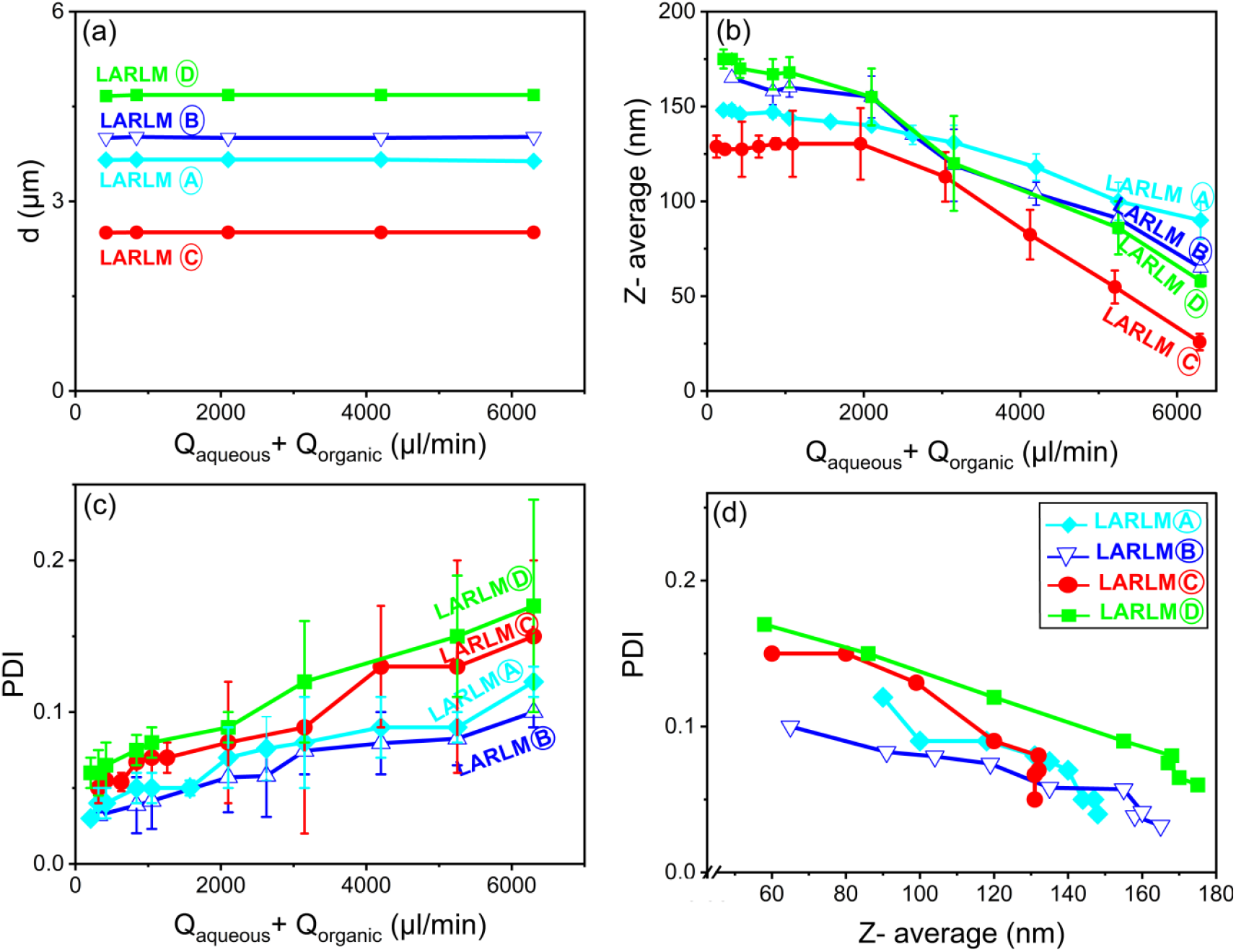
(a) Solvent sheet thickness *d* at 9000 µm downstream from the injection point in dependence of *Q*_*aqueous*_ + *Q*_*organic*_ as obtained in simulations for *Q*_*aqueous*_ /*Q*_*organic*_= 20. (b) Experimentally obtained average size in dependence of *Q*_*aqueous*_ + *Q*_*organic*_ . (c) PDI of LNP measured with DLS for *Q*_*aqueous*_ /*Q*_*organic*_ = 20 in dependence of *Q*_*aqueous*_ + *Q*_*organic*_ . (d) PDI in dependence of Z-average. (From left to right the flow rate is decreasing and data are generated for (*Q*_*aqueous*_ /*Q*_*organic*_ = 20). Error bars represent standard deviations obtained in 9 experiments (3 repetitions for each fabricated device, 3 devices were fabricated for each design). No error bars are given in (d) but can be taken from (b) and (c) for particles size and PDI respectively.

The production of smaller nanoparticles seems always to be coupled with some increase in PDI. Moreover, the desired smallness of the nanoparticles can be achieved under adjusted operation conditions with all LARLM designs. Therefore, the performance of the different LARLM could be best compared based on the dependence of the PDI on the particle size as shown in figure 5d. LARLM Ⓑ demonstrated the lowest PDI values among all LARLMs tested. The PDI values for LARLM Ⓑ remained below 0.1 across the particle size range, indicating highly monodisperse populations meeting the quality standards for pharmaceutical applications. LARLM Ⓑ provides a balanced environment for nucleation and growth processes, minimizing the stochastic variations that typically lead to broader size distributions. A reduction of z-dimensions as in LARLM Ⓓ that leads further away from the ideal of one-dimensional flow (infinite z dimension) seems to compromise the performance. To our surprise, the narrower injection nozzle of LARLM Ⓐ that in comparison to LARLM Ⓑ pushes the initial thickness of the organic phase closer to the final one seems not to improve the performance. The reduction of velocity differences between the two phases but also other effects like reduced homogeneity of the organic sheet layer thickness that become more influenced by fabrication tolerancesat reduced dimensions may play a role. In comparison to LARLM Ⓑ the reduction of the main channel cross section in *y*-direction as realized in LARLM Ⓒ compromises the performance as well. Probably the outer flow of aqueous phase become more affected by small fabrication tolerances leading to inhomogeneities in the organic sheet thickness. In the following only LARLM Ⓑ giving also the lowest PDI values will be further evaluated. Comparing the designs and realizing that higher volume flows lead to a reduction in particle sizes requires a more in-depth discussion of the development of the flow profile behind the injection nozzle.

Figure 6a shows how a velocity in the centerline *v* (in the middle of the injected sheet) as obtained in simulations for LARLM Ⓑ develops in dependence of *x*-position. The flow velocity immediately after leaving the nozzle slows down initially, with this decrease being more pronounced at higher values for *Q*_*aqueous*_ + *Q*_*organic*_ (*Q*_*aqueous*_ /*Q*_*organic*_ = const = 20). This decrease is associated with an expansion of the emerging organic phase and leads to greater shear between the organic and aqueous phases. (Figure 1a-d Supplementary Information). Then *v* quickly rises before reaching a plateau, indicating a fully developed flow (see Figure 7a). The point where *v* reaches 90% of the saturation value was defined as the stabilizing position *x*_*stablizing*_. With the Reynolds numbers calculated for the channel outlet (between 12.5 and 190 depending on the total flow rate) the entrance length as specified in the literature^22^ was calculated to be 170 µm to 1550 µm and corresponded quite well with *x*_*stablizing*_ . The diffusion time was calculated as 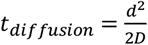 (*D* the constant for diffusion of ethanol in water) and ranged from 1.5 ms for LARLM Ⓒ to 10 ms for LARLM Ⓓ and the corresponding position *x*_*diffusion*_ at which diffusion is completed was derived as:

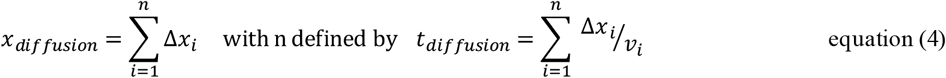

where *v*_*i*_ was the velocity in the finite element placed in the center positions in the injected sheet cross sections at *x*-position indexed with *i* and Δ*x*_*i*_ the axial length of this finite element. Notably, for all studied flow rates, it was confirmed that molecular diffusion completed shortly after injection and before *x*_*stabilizing*_ was reached. To convert the *v*(*t*) dependency in figure 6a into a *t*(*x*) dependency as shown in figure 6b we used again the velocity information in the finite elements as:

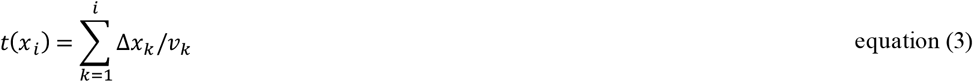

Moreover, the stabilizing time *t*_*Stablizing*_was determined as

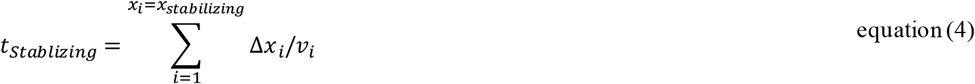

together with *t*_*diffusion*_ also in figure 6b and it shows that the except for the lowest flow rate that the diffusion is finished always *1*.*1 ms* before the flow profile has stabilized.

**Figure 6.**
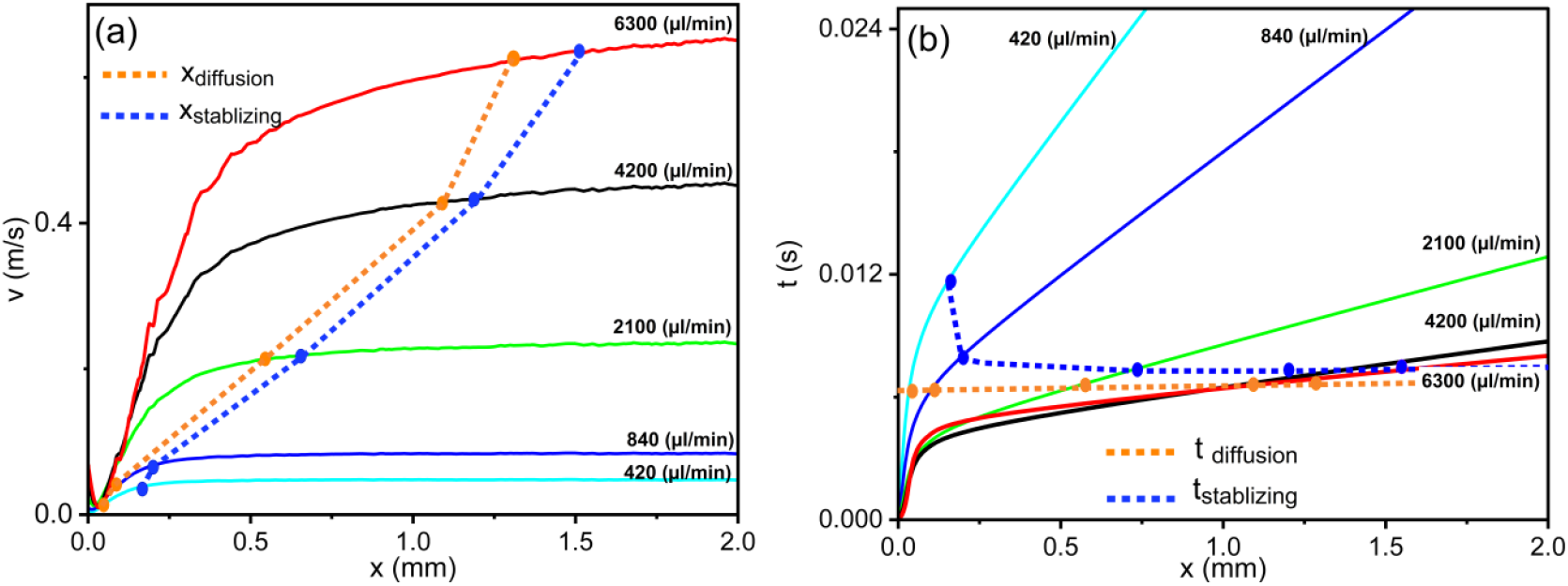
Analysis of the regimes of flow stabilization and diffusional mixing in LARLM Ⓑ at fixed flow rate ratio *Q*_*aqueous*_ /*Q*_*organic*_ = 20 and varied values of *Q*_*aqueous*_ + *Q*_*organic*_ . (a) Velocity profile *v*(*x*) along the channel centerline (*x*-direction) from the nozzle outlet to 4 mm downstream for total flow rates ranging from 420 µL/min to 6,300 µL/min. The *x*_*stablizing*_ and diffusion completion *x*_*diffusion*_ are indicated by dashed vertical lines, with their intersection points on the velocity curves highlighted. The initial drop in velocity in the range up to 200 µm downstream of the nozzle is examined in more detail in Supplementary Information 1. (b) Profile of elapsed time *t*(*x*). The dashed lines represent the characteristic timescales for *t*_*Stablizing*_ and *t*_*diffusion*_ .

**Figure 7.**
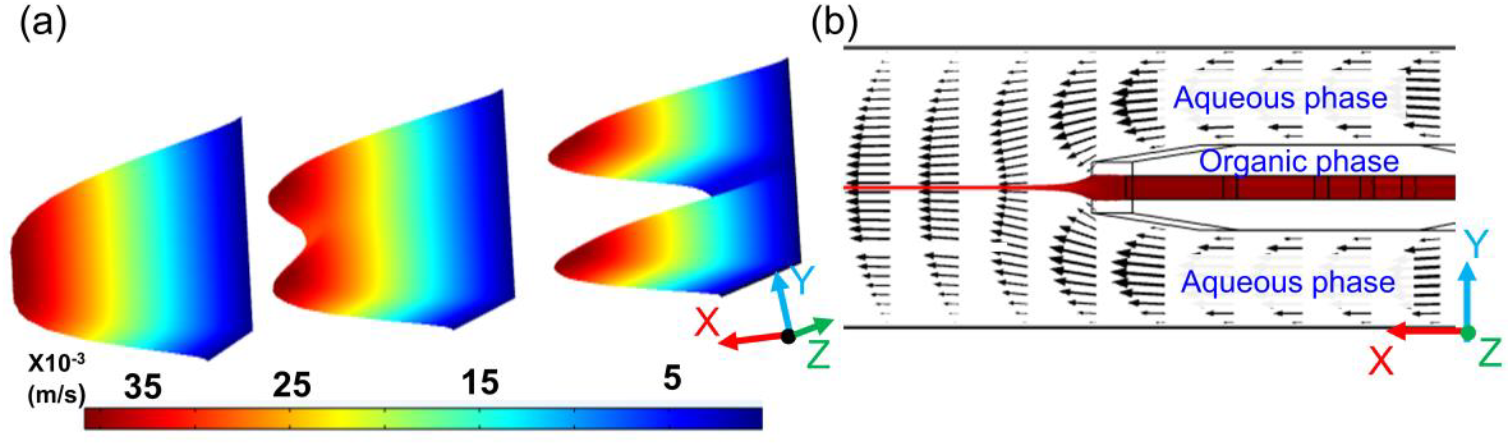
(a) 3D velocity profiles in the channel at critical downstream locations (x = 30, 100, 200 µm), revealing progression from initial instability to steady-state flow. (b) Simulated velocity field showing streamlines in the LARLM Ⓑ viewed as a cut in the x-y plane, where the length of the arrows is proportional to the local velocity ( *Q*_*aqueous*_ + *Q*_*organic*_ = 420 μL/min).

These results demonstrate that diffusion is completed within a relatively short distance from the nozzle, while flow stabilization extends further downstream. Up to flow rates of 2000 µL/min, the particle sizes did not vary. As total flow rates further increase, *x*_*diffusion*_shifts downstream, but *xSt*_*ablizing*_ even more. When the flow stabilization shifts downstream this has an influence on the growth of precipitated particles. As illustrated by the simulations shown in figure 7a, b the organic phase, which is squeezed into that thin central sheet, moves more slowly than the aqueous phase as long as the flow is not yet stabilized.

The influence on diffusion that can affect particle growth can be explained by an interfacial dispersion model (IDM) as sketched in figure 8. Similar to the well-known Taylor-Aries dispersion the interplay of radial diffusion with a profile of axial velocity leads to intensification of diffusion. The continuous velocity difference generates axial velocity gradients across the interface (see figure 8). The faster moving aqueous phase is constantly bringing fresh water (lower ethanol concentration) to the interface which increases the concentration gradient and speeds up the diffusion. The latter exchanges ethanol with water and thereby slows down the nanoparticle growth.

**Figure 8.**
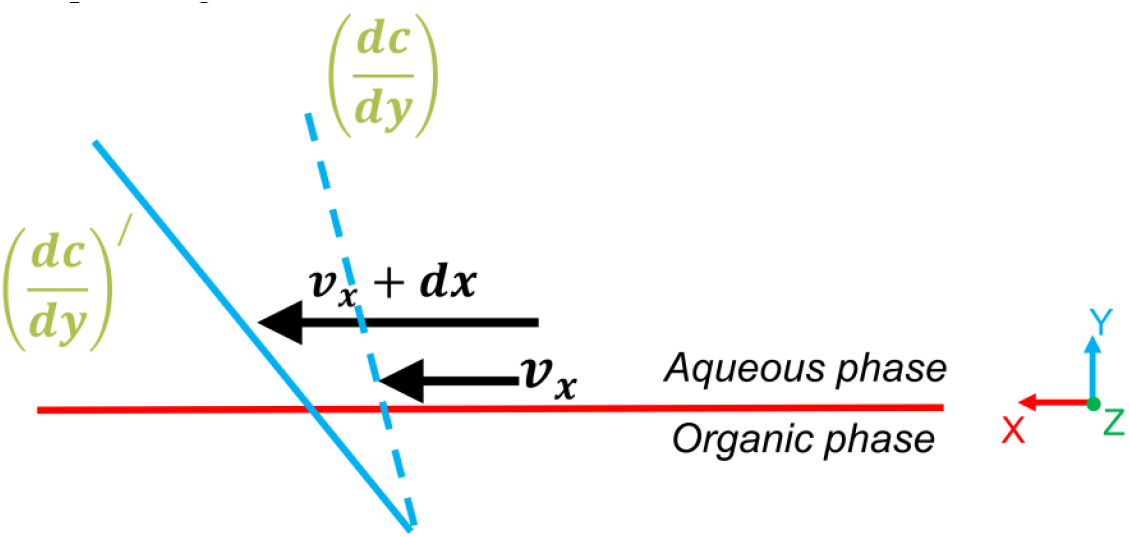
Sketch illustrating the IDM that leads to intensified diffusion as long as the flow has not stabilized. The concentration gradient in y-direction is thereby not only influenced by diffusion in y-direction but also by convection in x-direction.

For pharmaceutical formulation the consistency between batches and regulatory compliance are paramount, but also the production efficiency (throughput) must be sufficient to deliver samples in sufficient volumes to allow testing. The scalability of LNP production is demonstrated in figure 9, where in dependence of total flow rates the time required for 1 liter of LNP suspension is given together with LNP quality parameters. At 6300 µl/min, monodisperse 60nm nanoparticles are collected in only 2.65 hours. This represents a more than 15-fold time reduction compared to lower flow rates, while still maintaining perfect particle characteristics (Z-average = 60nm, PDI ≤ 1 with LARLM Ⓑ).

**Figure 9.**
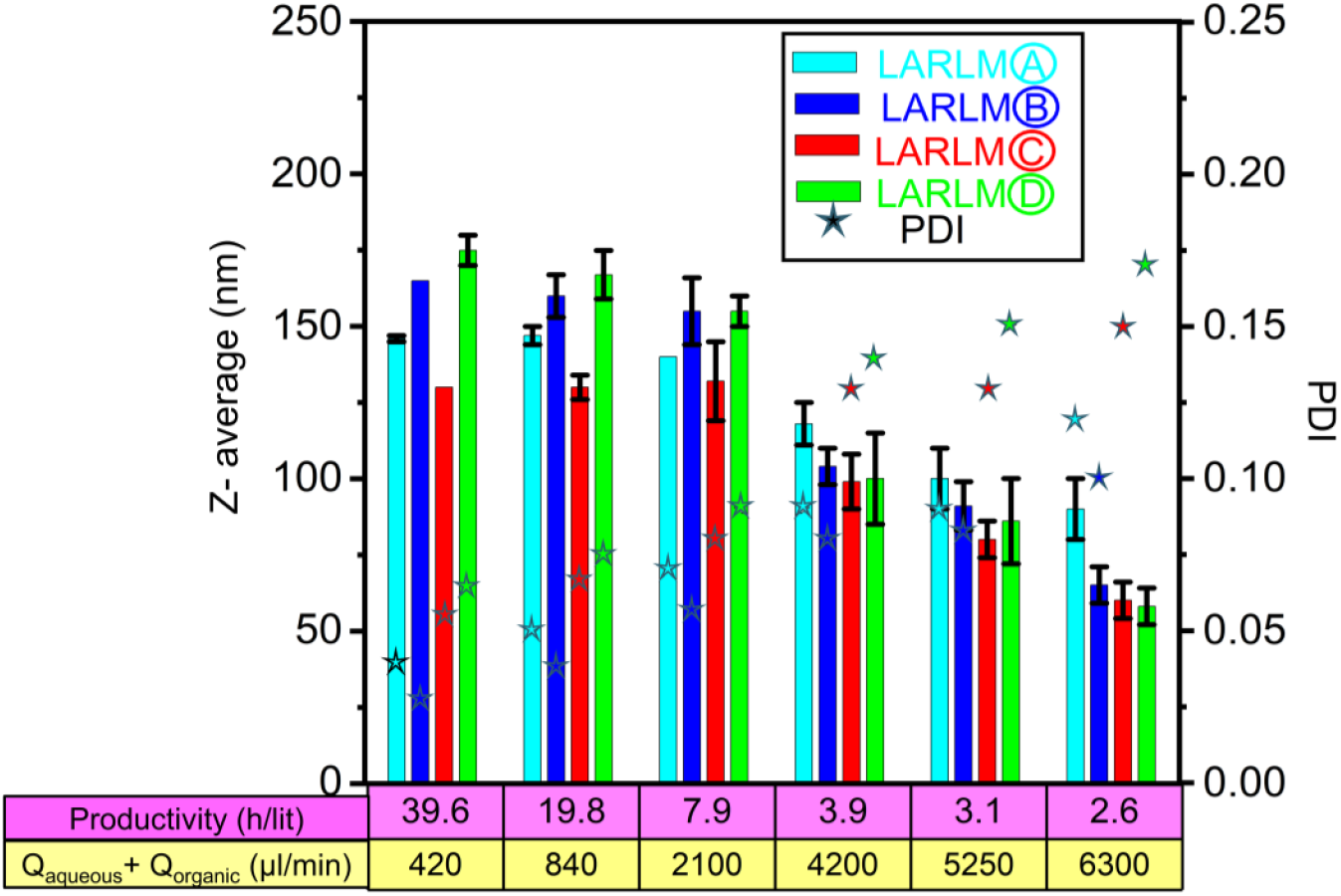
(a) Rapid high-volume nanoparticle production. Time required for 1L collection of 60nm LNPs at increasing total flow rates (*Q*_*aqueous*_ + *Q*_*organic*_ ). At 6300 µl/min (matching volumetric flow to collection scale), monodisperse particles (PDI < 0.1) are achieved in just 2.65 hours in LARLM Ⓑ.

Since LARLM Ⓑ gives the best option, it was used in the following analysis of encapsulation efficiency. LNP were loaded with the coumarine-6, simvastatin and diclofenac sodium salt in LARLM Ⓑ. Figure 10a shows that drug loading on the nanoparticle size. The particle size is only slightly increased at a higher total flow rate of *Q*_*aqueous*_ + *Q*_*organic*_ = 2100 µl/min. The PDI also indicates an almost monodisperse distribution, but the LNP loaded with simvastatin is higher than 0.1 at a flow rate of *Q*_*aqueous*_ + *Q*_*organic*_ = 4500 µl/min (See figure 10b). As it is shown in figure 10c the effective encapsulation of coumarin-6 within the lipid matrix will increase its concentration in the supernatant *C*_*en*_ after centrifugation, because free coumarin-6 precipitates and sediments in aqueous environment. The calibration curve (absorbance at 447 nm against known concentrations of coumarin-6) was used to quantify *C*_*en*_. Based on equ. (1). EE% was determined to be 71± 5% (at *Q*_*aqueous*_ /*Q*_*organic*_ = 20). While higher total flow rates can reduce LNP size, a variation of total flow rates (*Q*_*aqueous*_ + *Q*_*organic*_ = 420 – 6300 µl/min) did not alter the EE%

**Figure 10.**
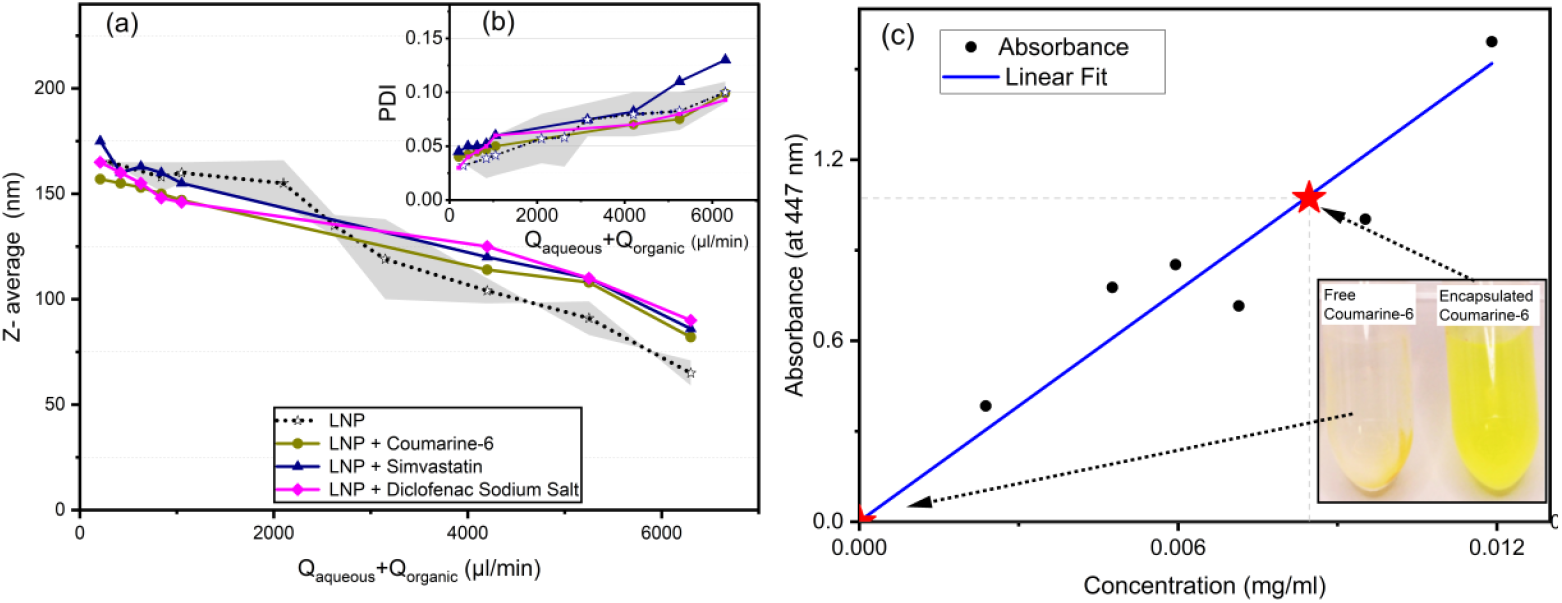
(a) Particle size and (b) PDI measured without drug encapsulated and with encapsulated coumarin-6, Simvastatin and Diclofenac sodium salt after LARLM processing at *Q*_*aqueous*_ /*Q*_*organic*_ = 20 and *Q*_*aqueous*_ + *Q*_*organic*_ = 420 *to* 6300 μL min^-1^. (c) Light absorption of the coumarin-6 measured in the supernatant after centrifugation of suspensions obtained by LARM processing at *Q*_*aqueous*_ /*Q*_*organic*_ = 20 and *Q*_*aqueous*_ + *Q*_*organic*_ =1050 μL min-1. The upper red star marks concentration *C*_*en*_ obtained on the linear calibration curve, which was offset-corrected by subtracting the absorbance in the supernatant of free coumarin-6 (identical microfluidic conditions but without lipid). The latter defines the zero point (red star) of the calibration curve. Two photos of vials illustrate the respective physical separation of coumarin-6 with and without lipid encapsulation.

To assess cellular uptake and persistence of LNPs generated by different flow rates *Q*_*aqueous*_ + *Q*_*organic*_ = 420 – 6300 µl/min, coumarin-6-loaded particles were tested in primary human endothelial cells. The encapsulation efficiency and particle size per condition (*Q*_*aqueous*_ + *Q*_*organic*_) were: 420 μL /min: 76% and 163 nm; 4200 μL min^-1^: 72% and 104 nm; 6300 μL min^-1^: 70% and 61 nm. Non-encaspulated coumarin-6 served as control. Cells were treated for 30 min after which intracellular green fluorescence was observed (see figure 11a). The signal was present after 4h and the cells did not show any morphological changes or signs of cell death.

**Figure 11.**
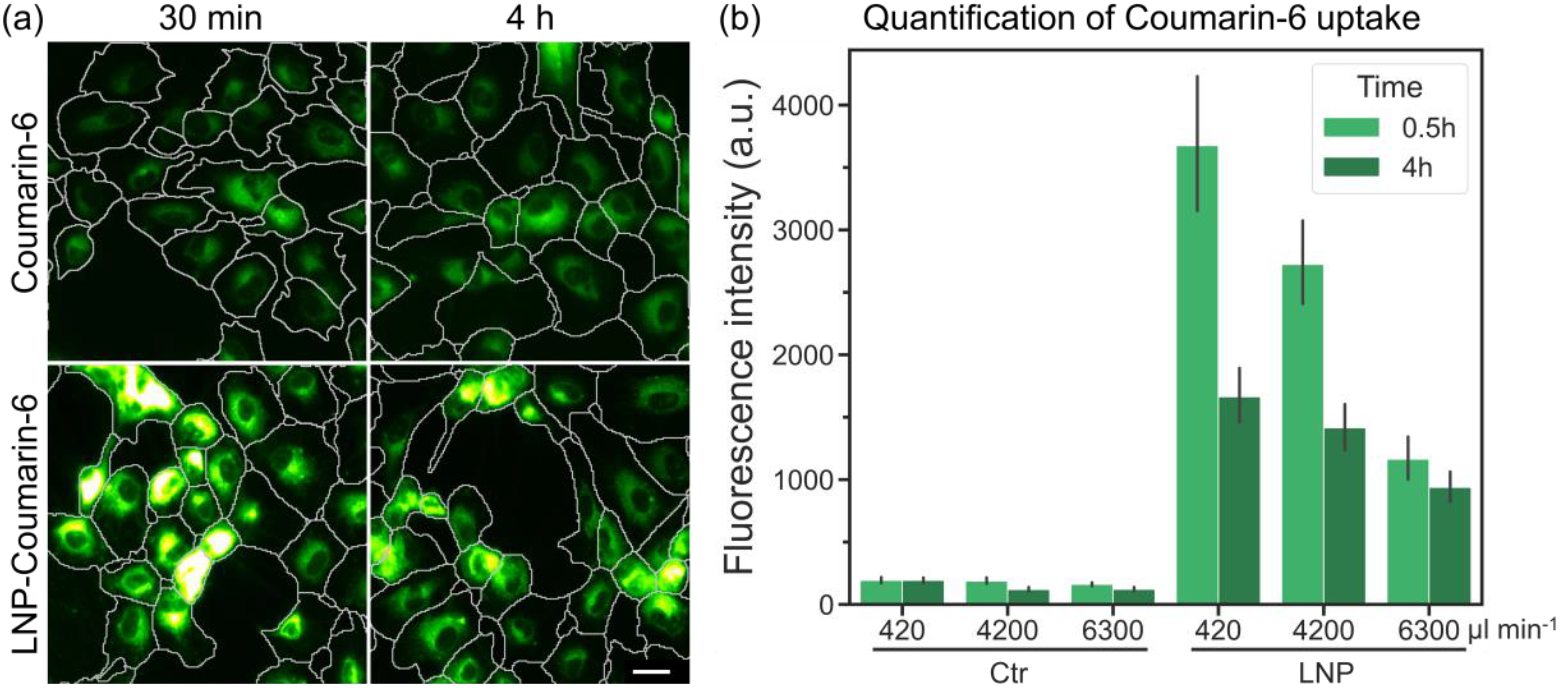
Cellular uptake of coumarin-6 with and without LNP encapsulation, generated by LARLM Ⓑ. a) LNP-treated cells showed increased coumarin-6 signal intensity compared to non-encapsulated coumarin-6 treated cells. Scale bar is 25 µm. b) Quantification of coumarin-6 signal intensity over time. The numbers below the bar graph indicate the total flow rates (*Q*_*aqueous*_ + *Q*_*organic*_) for control and LNP generation. Overall, cellular uptake and persistence of coumarin-6 was higher when administered using LNPs. Error bars indicate the 95% confidence interval, n = 3 biological replicates.

Quantification of coumarin-6 signal intensity revealed up to a 15-fold increase in uptake of coumarin-6 encapsulated in LNP compared to controls with free coumarin-6 (see figure 11b). Interestingly, while EE% was comparable for each condition, LNP samples generated at lower flow rates resulted in higher intracellular signal intensities than those generated at higher flow rates. While signal intensity decreased over time, it remained elevated in LNP-treated cells compared to controls. Notably, signal persistence was better for the LNP generated at high flow rates. In line with literature, larger LNPs may be preferentially internalized via endocytosis but are more prone to degradation or exocytosis^23^. Smaller LNPs, although sometimes taken up less efficiently, tend to persist longer intracellularly, potentially due to reduced lysosomal processing. These findings suggest a trade-off between the efficiency of cellular uptake and intracellular persistence based on LNP size, which is crucial for designing effective nanoparticle-based delivery systems for potential later drug-treatment.

## 4. Conclusion

Through the sophisticated fabrication enabled by 2PP, we engineered and evaluated several LARLM designs with varying channel and nozzle geometries. Our results confirm that the core principle behind all LARLMs, which is hydrodynamic focusing of the organic phase into a thin, central sheet within a co-flowing aqueous phase effectively addresses critical limitations of conventional microfluidics. Not only can contamination be avoided and long-term operation enabled, but throughput could also be increased to a level that has not been achieved with other microsystems before. An important finding was the identification of the design variant that delivered significantly better results and consistently produced LNPs with remarkably low polydispersity (PDI < 0.1) across all productivity levels and a wide range of particle sizes. In addition, we discovered a phenomenon that cannot be explained intuitively and was previously unknown, namely that an increase in the total flow rate above 2000 µL/min led to an unexpected significant reduction in particle size. This observation could be explained by a qualitative interfacial dispersion model (IDM) that takes into account the increased diffusive mixing caused by velocity gradients at the organic-aqueous interface before the flow profile stabilizes. Scalable production with a LARLM system was demonstrated by achieving a production rate of 1 liter of monodisperse (LNP size 75 nm) suspension in just 2.65 hours - a level of productivity that meets the volume requirements for preclinical and clinical studies. The versatility of the platform was further demonstrated by the successful encapsulation of various active ingredients, including coumarin-6, simvastatin, and diclofenac, with a high and consistent encapsulation efficiency (∼71%) for coumarin-6. Finally, in vitro studies on HUVECs provided compelling biological validation. LNPs produced by the LARLM significantly enhanced the cellular uptake and intracellular persistenceof coumarin-6 compared to its free form, with an observed inverse relationship between particle size (controlled by total flow rate) and signal persistence. This underscores the potential of our technology to tailor LNP properties for specific therapeutic outcomes, such as balancing uptake efficiency with intracellular retention.

In summary, the LARLM platform represents a significant advancement in microfluidic nanoparticle synthesis. It combines the precision of lab-scale devices with the throughput required for pharmaceutical manufacturing, all while ensuring operational stability and superior product quality. Since an earlier study has already shown that the LARLM design also enables in-situ measurement of particle size, this approach proves ideal for use in quality-assured and effective industrial production of LNP and, in particular, for supporting preclinical studies.

## AUTHOR INFORMATION

### Author Contributions

*Ebrahim Taiedinejad*

- Conceptualization
- Investigation
- Methodology
- Visualization
- Writing – original draft
- Writing – review & editing

*Alshaimaa Abdellatif*

- Methodology
- Visualization

*Victor Krajka*

- Conceptualization
- Visualization

*Andreas Dietzel*

- Conceptualization
- Supervision
- Writing – review & editing

## Conflicts of interest

There are no conflicts to declare for all authors/coauthors:

*Ebrahim Taiedinejad*

*Alshaimaa Abdellatif*

*Victor Krajka*

*Andreas Dietzel*

## Acknowledgement

The work was financially supported by Deutsche Forschungsgemeinschaft (DFG) and Fraunhofer Gesellschaft with funds for the trilateral transfer project,,DLS-Feedback-regulierte kontinuierliche Partikelproduktion” under grant number 426328385.

