## Supplementary information 1 for "Monodisperse LNPs - from Efficient Microfluidic Production and Loading to in Vitro Testing"

The simulations shown in Figure 1 illustrate how the organic jet exits the narrow nozzle and enters the wider mixing zone, where it meets the aqueous phase. A strong expansion of the jet can be observed directly at the nozzle outlet. According to the continuity equation, the expansion leads to a reduction in velocity. Therefore, the velocity initially drops sharply after the nozzle before rising again. At At $Q_{aqueous}/Q_{organic} = 20$, the aqueous phase streams initially move faster than the organic jet on both sides, creating a steep velocity gradient at the interface. The simulations show that this gradient is even more pronounced at higher total flow rates.

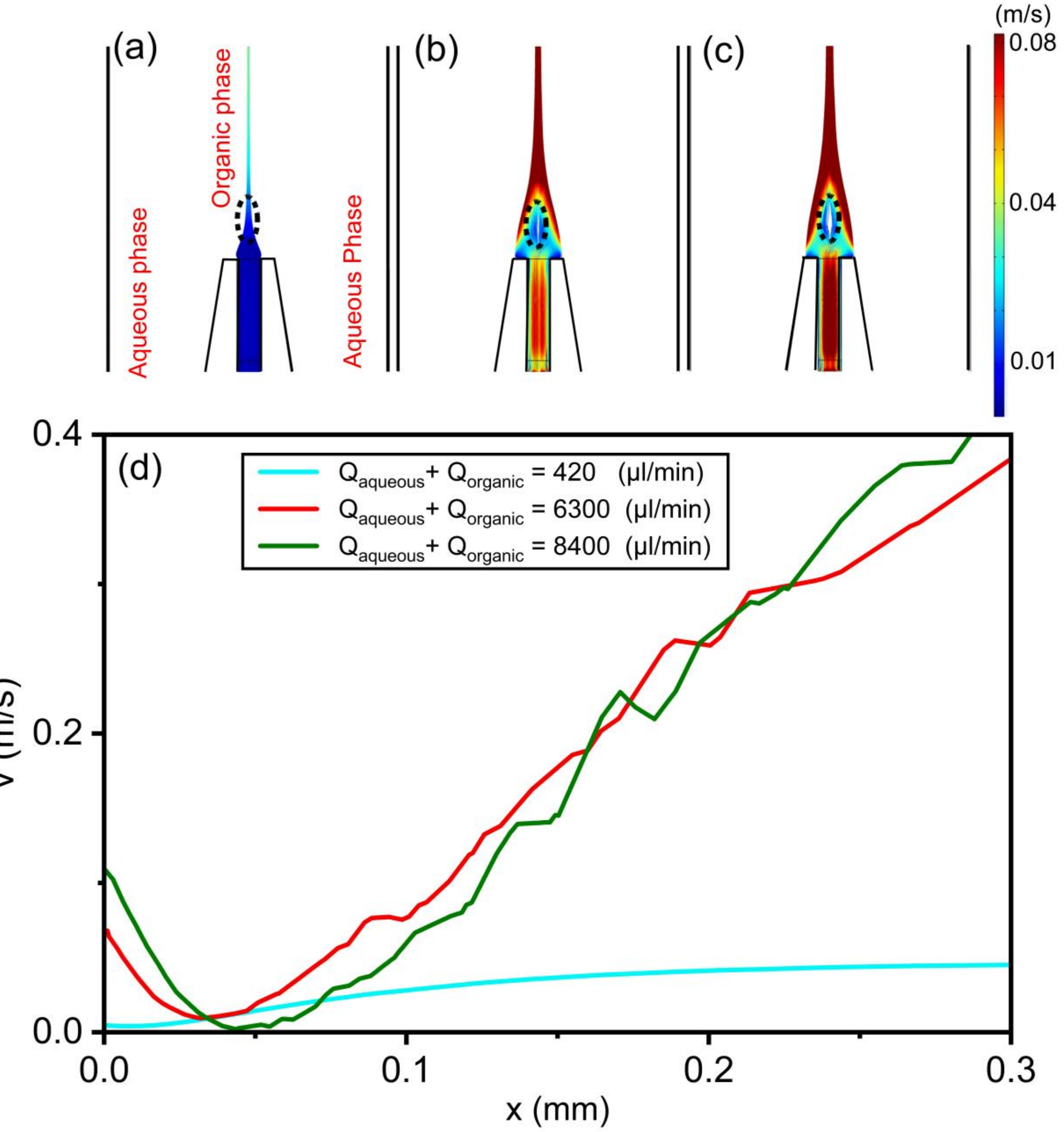


Figure 1: Simulated velocity field (x/y cross-sectional view) obtained at a flow rate ratio of $Q_{aqueous}/Q_{organic}$ = 20 at the nozzle in the LARLM Ⓑ for $Q_{aqueous} + Q_{organic}$ = (a) 420 (µL/min), (b) 6300 (µL/min), and (c) 8400 (µL/min). The area outlined in dashed lines indicates where the velocity initially decreases downstream of the nozzle. (d) The axial centerline velocity close to the nozzle also shows the initial velocity drop , which becomes more pronounced at higher total flow rates.
